# The best of both worlds: A new lipid complex has micelle and bicelle-like properties

**DOI:** 10.1101/437327

**Authors:** Monica D. Rieth

## Abstract

Bicelles have been demonstrated to be a valuable tool for studying membrane protein interactions and structure *in vitro*. They are distinguished by a distinct lipid bilayer that mimics the plasma membrane of cells making it more native-like than its detergent micelle counter-part. Bicelles are typically comprised of a long-chain phospholipid such as dimyristoylphosphatidylcholine (DMPC) and a short-chain phospholipid such as dihexanoylphosphatidylcholine (DHPC). When mixed together in solution DMPC-DHPC bicelles assume a discoidal structure comprised of a heterogeneous arrangement where the short-chain lipids gather around the rim of the disk and the long-chain lipids form the flat, planar, bilayer region. In this study, the nonionic surfactant, C_8_E_5_, was used to prepare mixtures with DMPC to determine if it adopts properties similar to bicelles with a ***q*** ≥ 0.5. At ***q*** ≥ 0.5, DMPC-DHPC bicelles are bilayered and DMPC is sequestered from the detergent micelle-like DHPC. Mixtures of DMPC and C_8_E_5_ were prepared at various ***q*** values, a parameter used to describe the mole ratio of DMPC to DHPC in the preparation of bicelles. Employing biophysical methods like dynamic light scattering, ^31^P-NMR and analytical ultracentrifugation, properties of these lipid-detergent complexes are described. Interestingly they adopted a spherical-shaped micellar structure morphology and did not assume a discoidal shape typical of bicelles at ***q*** ≥ 0.5. However, they appear to retain bilayer-like properties that may prove beneficial for *in vitro* biophysical studies of membrane proteins.

## 1. Introduction

It has been well-established that 1,2-dimyristoyl-*sn*-glycero-3-phosphocholine (DMPC), a naturally derived phospholipid with a 14-carbon chain length and a choline head group, spontaneously assembles into discoidal lipid structures called bicelles when mixed with short-chain phospholipids such as 1,2-dihexanoyl-*sn*-glycero-3-phosphocholine (DHPC) - also naturally derived but containing a 6-carbon chain length with a choline head group [1, 2]. These bilayered structures can also be prepared using DMPC and bile salt detergents such as CHAPS or CHAPSO in lieu of a short-chain phospholipid [1, 3, 4]. DMPC-DHPC bicelles are among the most extensively characterized bicellar structures, and they have been used successfully as membrane mimics in solution NMR studies of membrane protein structure [4 – 7]. These lipid complexes can be tailored to different applications by altering the properties of the lipids used in their preparation. For example, the lipid tail length and the ratio of DMPC to DHPC can affect the resulting physical properties of bicelles. Anionic lipids can also be included to alter the charge at the surface [8]. Collectively, these properties affect the thickness and the planar length of the bicellar structure both of which can influence membrane protein structure when they are incorporated and ultimately, their stability [9]. Bicelles are of unique interest for membrane protein studies because unlike their detergent counterpart, they contain a bilayered planar region, which captures a true membrane-like environment (Figure 1) [4, 5]. In membrane protein studies, choosing the optimal detergent or lipid system usually necessitates a screening of potential candidates because not all systems are universally adaptable to all types of membrane proteins [10 – 13]. In this study, a novel lipid-detergent complex was prepared with DMPC, the long-chain phospholipid, and a nonionic detergent, pentaethylene glycol n-octyl ether or C_8_E_5_ (Figure 1). Physical properties of the resulting complexes were characterized to assess whether bicelle-like properties were retained. Two parameters commonly used to describe bicelles were investigated. The ***q***-value, which describes the ratio of the long-chain lipid to that of the short-chain lipid and the total lipid concentration (% w/w), which is defined as the total concentration of DMPC + C_8_E_5_. Our analysis shows the lipid complexes assume a shape that is closer to spherical than discoidal. Further, at various ***q***-values, DMPC-C_8_E_5_ complexes show remarkable changes in solution viscosity that are not typically observed for DMPC-DHPC bicelles at ***q*** ≥ 0.5. Because of this, the ***q***-value for DMPC-C_8_E_5_ samples was kept at a ***q*** ≤ 0.5, a range we refer to as the “solution dynamic range.” All ***q***-values reported were determined on a theoretical basis and do not account for any free DHPC and C_8_E_5_ detergent in solution. It should be acknowledged that accounting for free detergent in solution based on the critical micelle concentration results in a larger observed ***q, q***_eff_ [2]. For example, ***q*** = 0.7 bicelles, results in a ***q***_eff_ = 1.0 determined from the CBC of DHPC [2, 14]. This accounts for the DHPC associated with bicelles, which is in equilibrium with free DHPC in solution.

**Figure 1.**
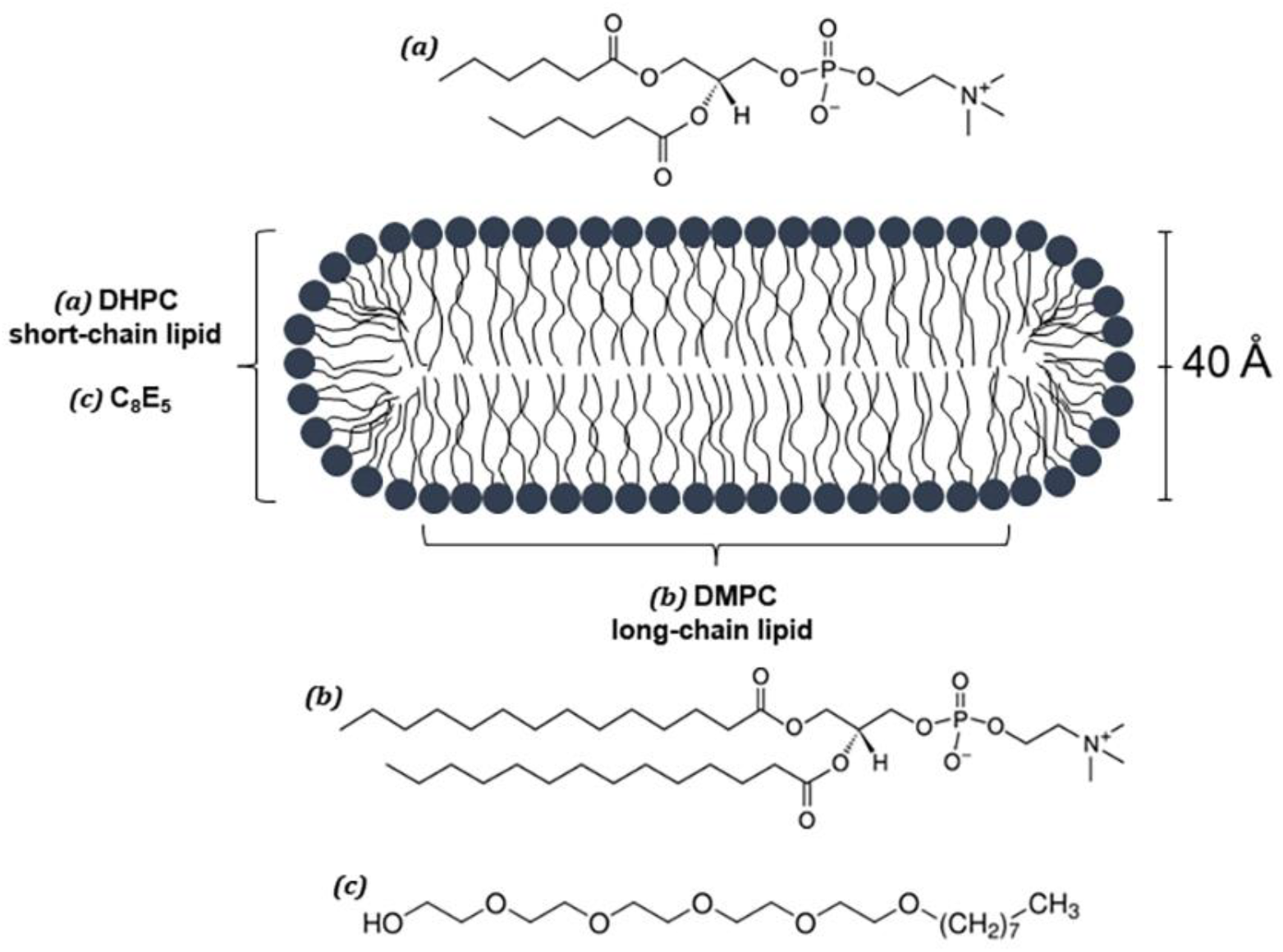
Schematic diagram of a ***q*** ≥ 0.5 bicelle containing DMPC and DHPC. Bicelles with a ***q*** ≥ 0.5 have an appreciable bilayer region. C_8_E_5_ lipid complex samples at ***q*** ≤ 0.5 contain the C_8_E_5_ detergent on the rim instead of DHPC.

**Figure 2.**
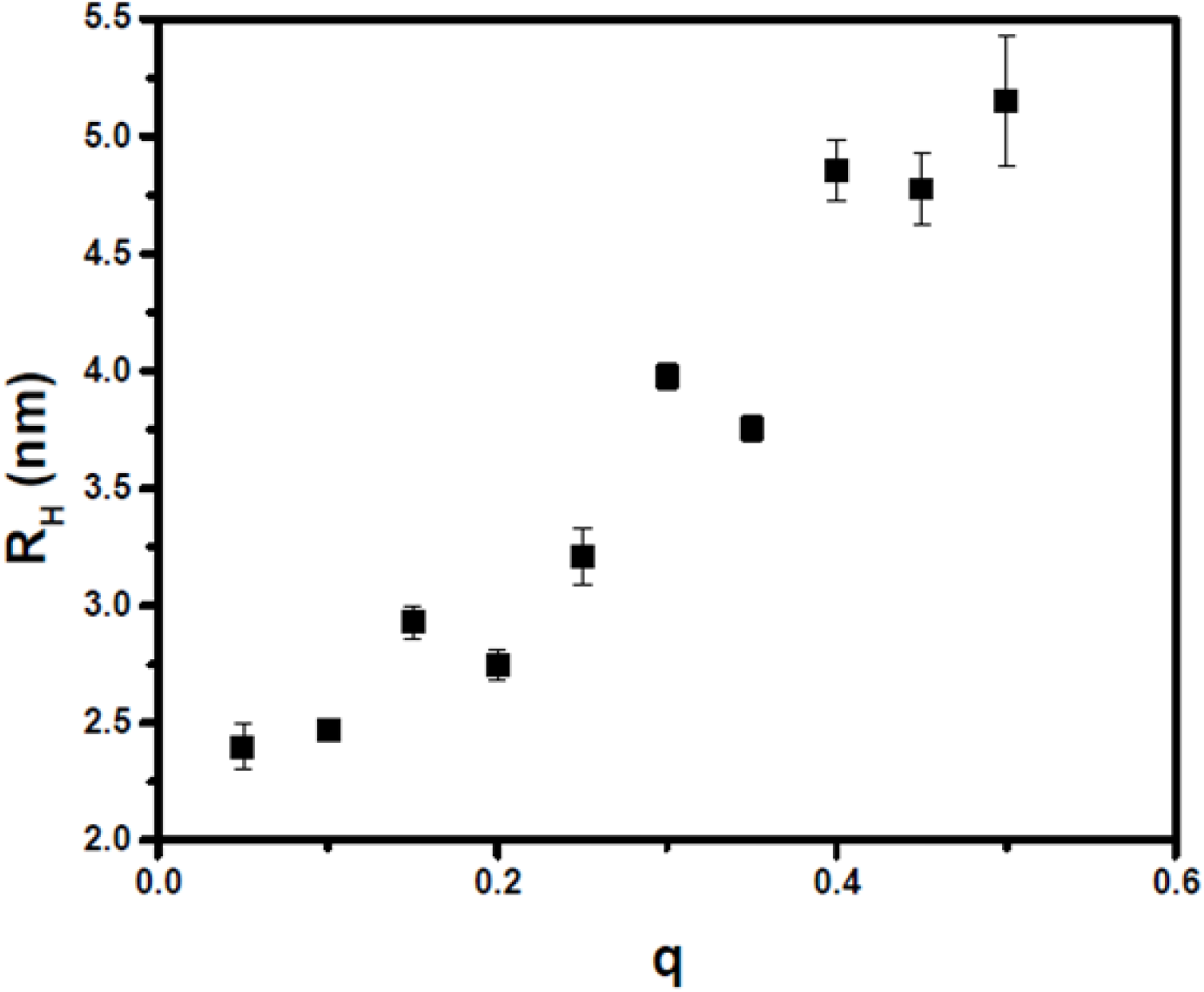
Dynamic light scattering data measuring R_h_ as a function of ***q***. Increasing ***q*** corresponds to an increase in the hydrodynamic radius of the particle for DMPC-C_8_E_5_ complexes. All ***q*** values were kept below 0.5.

**Figure 3.**
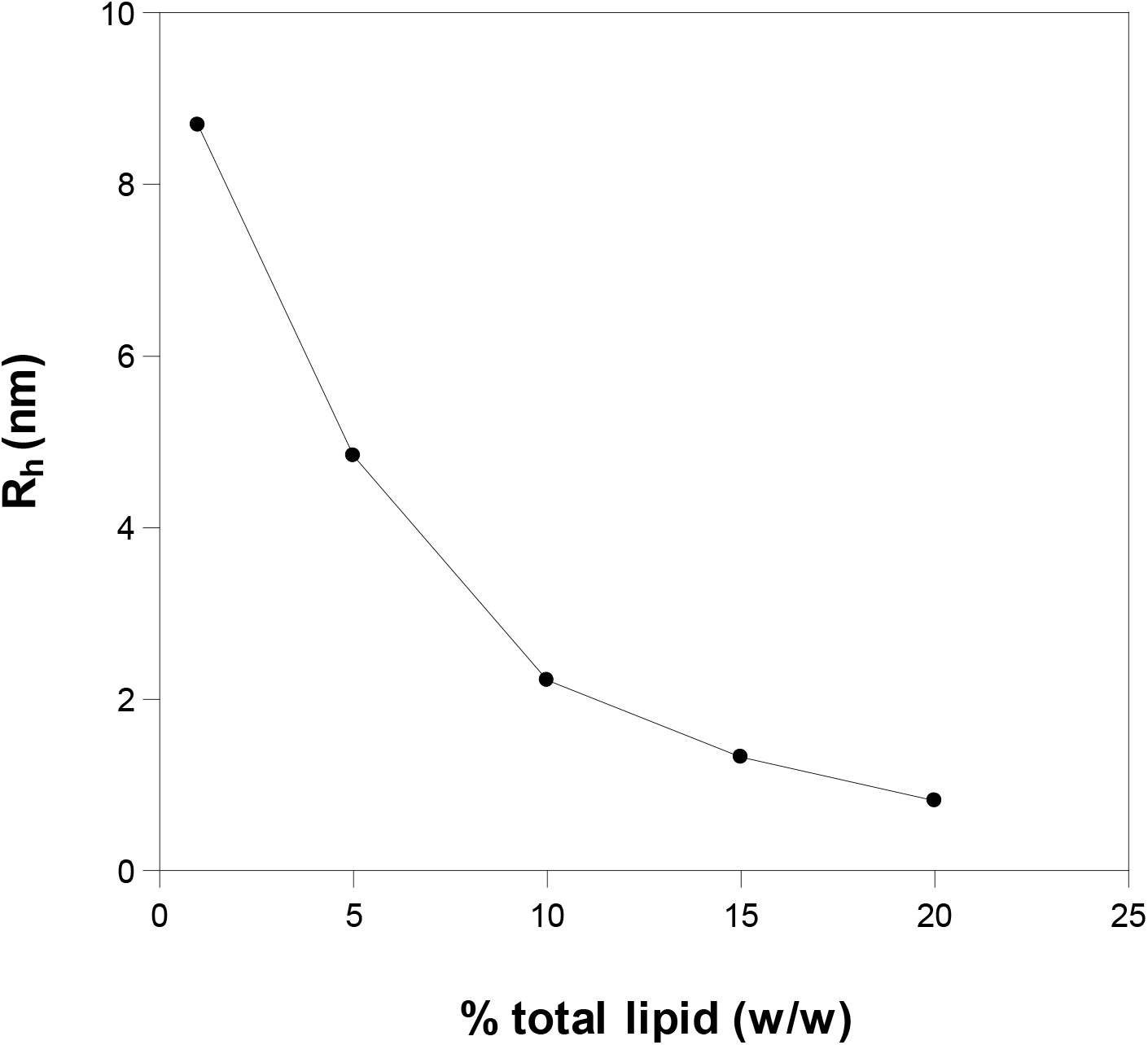
Dynamic light scattering data measuring R_h_ as a function of % total lipid (w/w). All samples were prepared at a ***q*** = 0.25. Particle size decreases as total lipid concentration increases.

In addition to characterizing important parameters of these complexes, we also show that they can be adapted to studies of membrane protein self-association (oligomerization) using sedimentation equilibrium in the analytical ultracentrifuge. A method that has been used successfully to investigate membrane protein oligomerization in both micellar and bicellar environments [2, 15]. With a partial specific volume similar to water, 0.983 cm^3^ / g, C_8_E_5_ micelles require little need for density matching [16]. However, in lipid complexes, only a small quantity of the density modifier deuterium oxide, D_2_O, was required for matching.

## Materials and methods

### 2.1 Chemicals and reagents

DMPC (1,2-dimyristoyl-*sn*-glycero-3-phosphocholine) and NBD-DMPE (1,2-dimyristoyl-*sn*-glycero-3-phosphoethanolamine-N-(7-nitro-2-1,3-benzoxadiazol-4-yl) (ammonium salt)) were purchased from Avanti Polar Lipids (Alabaster, AL, USA). C_8_E_5_ (n-octylpentaoxyethylene) was purchased from Bachem (King of Prussia, PA, USA). All lipids and detergents were used without further purification. NBD-DMPE was incorporated in small molar quantities so that absorption optics could be used to analyze the lipid complexes in the analytical ultracentrifuge. HEPES (2-[4-(2-hydroxyethyl)piperazin-1-yl] ethanesulfonic acid) was purchased from EMD Millipore (Billerica, MA USA) and NaCl (sodium chloride) was purchased from Avantor (Center Valley, PA, USA). D_2_O (deuterium oxide) was purchased from Cambridge Isotopes (Tewksbury, MA, USA). Milli-Q water (Millipore) was used throughout all experiments.

### 2.2 Preparation of DMPC-C_8_E_5_ mixtures

#### 2.1.1 Samples for analytical ultracentrifugation

DMPC-C_8_E_5_ lipid solutions were prepared and evaluated at ***q*** = 0.25 and a total lipid composition of 5% (w/w). A small amount of NBD-labeled DMPE was incorporated into DMPC-C_8_E_5_ samples to allow the lipid complexes to be monitored using absorption optics. The ratio of NBD-labeled lipid to unlabeled lipid was kept sufficiently low so that the label did not influence the physical properties of the complexes (1:500 mole ratio of NBD-DMPE:DMPC). To a 1.5 mL eppendorf tube a sufficient quantity of 50 mg / mL DMPC in chloroform was added to give the appropriate ***q*** value. An appropriate quantity of 66 μg / μL of NBD-DMPE chloroform solution was immediately added to yield a 1:500 mole ratio of fluorescent-labeled lipid to unlabeled lipid. The samples were briefly vortexed and spun in a microfuge. 500 μL of Millipore water was added immediately before freezing the samples in liquid nitrogen. The samples were lyophilized to a powder. To each lyophilized sample an appropriate amount of Millipore water was added followed by a 40X concentrated stock solution of buffer, which yielded a final concentration of 10 mM HEPES, 100 mM NaCl pH 7.4. The samples were vortexed to a milky white suspension and C_8_E_5_ detergent was added to each sample in quantities that would assume a ***q***-value of 0.25. Samples were vortexed for several seconds to clarity. D_2_O was added to each sample ranging from 0% (by volume) up to 25 % for a total of nine samples. Reference samples (buffer only) were prepared to a final volume of 150 μL according to the following: To a 1.5 mL eppendorf tube was added 146.0 μL of Millipore water (less D_2_O) followed by 40X concentrated buffer solution to final composition of 10 mM HEPES 100 mM NaCl pH 7.4.

Samples were loaded into a 6-channel equilibrium charcoal-filled epon centerpiece with a pathlength of 1.2 cm. 120 μL of each sample and reference was loaded in the appropriate channels. All sedimentation equilibrium studies were carried out on a Beckman XL-A analytical ultracentrifuge at 25 °C using a four-hole An-Ti 60 rotor. Absorbance spectroscopy was used to monitor the sedimentation of the lipid-detergent complexes down the solution column at a wavelength of 464 nm, which corresponds to the absorbance of the NBD-labeled lipid. Samples were initially equilibrated at 10,000 rpm (8,050 x g) for 16 hours to allow equilibrium to be reached. The point of equilibrium was evaluated at each speed using Match in the program, *HeteroAnalysis* (University of Connecticut, Storrs CT). Thereafter speeds were increased by 5,000 rpm up to 30,000 rpm (72,446 x g), and samples were left to equilibrate for 6 hours at each speed before acquiring data. Data was collected at rotor speeds of 15,000 rpm (18,112 x g), 20,000 (32,198 x g), 25,000 rpm (50,310 x g), and 30,000 rpm (72,446 x g). The sedimentation equilibrium profiles were analyzed and the data were fitted to the Lamm equation, Eq. 1, to extract the buoyant MW, M_eff_, of the lipid-detergent complexes in each of the three samples:

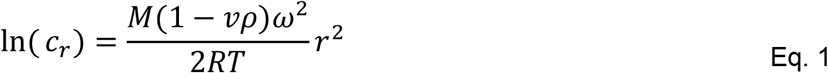

where *c*_*r*_ is the concentration of NBD-DMPE at radius *r, M* is the calculated molecular weight of the lipid-detergent complex, *v* is the partial specific volume of the lipid-detergent complex which was measured using a Kyoto density meter, *ρ* is the density of the HEPES buffer containing NaCl pH 7.4, which was also measured using the Kyoto density meter. *ω* is the angular velocity of the rotor, *r* is the radius from the center axis of rotation, *R* is the Boltzmann constant and *T* is the temperature in Kelvin.

#### 2.2.2 ^31^P-NMR sample preparation

Samples with a 25% (w/w) total lipid composition were prepared for ^31^P-phosphorus NMR experiments. Each sample was prepared at a ***q*** = 0.5 on a 990 μL scale. To two separate 1.5 mL eppendorf tubes, DMPC was added to a final concentration of 158 mM. To both samples 81.45 μL of Millipore water was added followed by 99.0 μL of D_2_O (10% (v/v)) and 6.6 μL 3.0 M sodium acetate pH 5.6 buffer. The samples were vortexed to a milky, white suspension. Next, 555.44 μL of a 25% (w/w) DHPC solution was added to one sample and 222.48 μL of a 50% C_8_E_5_ (w/w) solution was added to the second sample. The samples were vortexed to clarity. Last, praseodymium (III) chloride powder was added to a final concentration of 197 mM as per Glover et. al [4]. This concentration was previously determined to saturate the lanthanide-phospholipid interaction making the DMPC and DHPC ^31^P-phosphorus signals distinct from one another in the NMR [17].

NMR experiments were carried out on a Bruker DRX500 spectrometer equipped with a BBI probe. One dimensional ^31^P-phosphorus NMR spectra were recorded at 25 °C and 37 °C using a proton-decoupled single-pulse experiment with a minimum of 16 scans and a 100 ppm sweep width. The experiments were processed in TopSpin v. 1.3 (Bruker Corporation).

#### 2.2.3 Samples for DLS

Mixtures of DMPC-C_8_E_5_ were prepared based on the bicelle ***q***–value. ***q*** is determined by the mole ratio of DMPC to DHPC according to the following equation:

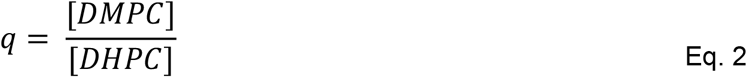

DMPC-C_8_E_5_ samples with the following ***q***-values were prepared: 0.05, 0.10, 0.15, 0.20, 0.30, 0.35, 0.40, 0.45, and 0.50 with a total lipid-detergent concentration kept constant at 5% (w/w). Samples were prepared on a 12 mL volume scale. To nine 15-mL centrifugal tubes, DMPC was added to achieve the bicelle ***q***-values described above. Next, 11.10 mL of Millipore water was added followed by 300 μL of 400 mM HEPES, 4.0 M NaCl pH 7.4 buffer. The DMPC mixture was vortexed for several seconds to create a milky, white suspension. To each of the samples, C_8_E_5_ was added in quantities that yielded ***q***-values of 0.05, 0.10, 0.15, 0.20, 0.25, 0.30, 0.35, 0.40, 0.45, and 0.50, respectively. The samples were vortexed after adding C_8_E_5_ and turned clear.

To determine the size dependence of DMPC-C_8_E_5_ lipid complexes on the total lipid-detergent concentration six samples were prepared on a 12 mL volume scale with a total lipid concentration of 1, 5, 10, 15, 20 and 25% (w/w). The ***q***-value was held constant at 0.25 for all six samples: To six 15 mL conical tubes DMPC was added to produce a final lipid-detergent concentration of 1, 5, 10, 15, 20, 25% (w/w) followed by the appropriate quantity of Millipore water. To each of the six samples 300 μL of 40X HEPES; NaCl pH 7.4 buffer was added to a final concentration of 10 mM HEPES; 100 mM NaCl pH 7.4 was also added to each sample. The samples were vortexed to a milky, white suspension and C_8_E_5_ was added in the proper amounts for a final lipid-detergent concentration of 1, 5, 10, 15, 20, and 25% (w/w) and a ***q***-value of 0.25. The samples were vortexed briefly until clear.

Viscosity measurements were carried out on a Cannon Fenske viscometer at a temperature of 20 °C and dynamic light scattering experiments were carried out on an ALV-CGS3 compact goniometer equipped with a 22 mW HeNe laser at a wavelength of 632.8 nm. The scattered light intensity was measured at a detector angle of 90 degrees for eight samples. Diffusion constants for the lipid-detergent complexes were acquired from the decay of the autocorrelation function, and the hydrodynamic radii were determined by the Stokes-Einstein equation for a spherical particle [18]. Results are summarized in Tables 1 and 2.

**Table 1.**
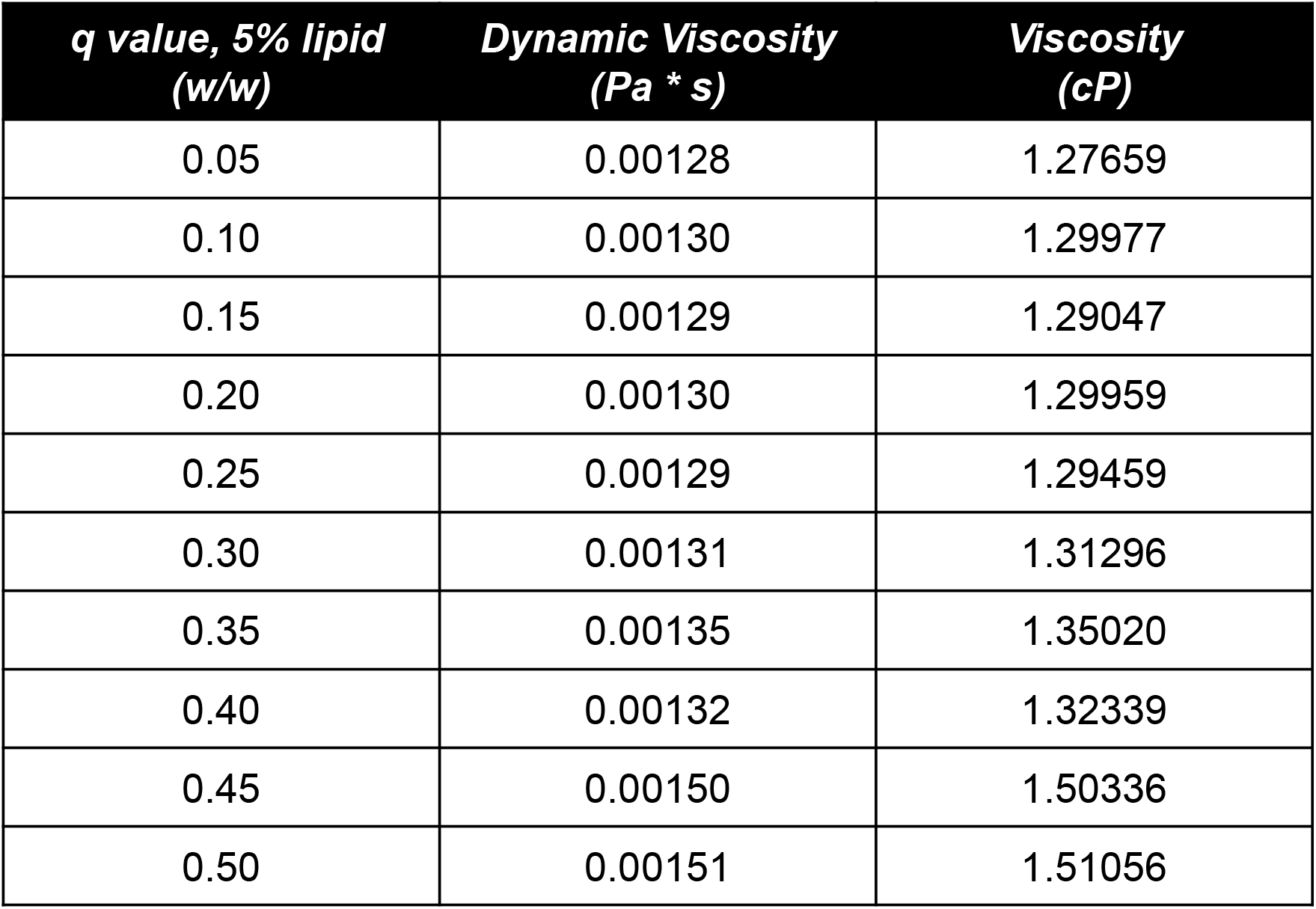
Viscosity changes as a function of ***q***. Viscosity increases with ***q***.

**Table 2.** Viscosity and R_h_ changes as a function of total lipid concentration.

| **$q = 0.25$,**<br>**% lipid (w/w)** | **Dynamic**<br>**Viscosity**<br>**(Pa\*s)** | **Viscosity**<br>**(cP)** | **$R_h$**<br>**(nm)** |
| --- | --- | --- | --- |
| 1 | 0.00109 | 1.09144 | 8.69 |
| 5 | 0.00129 | 1.28765 | 4.84 |
| 10 | 0.00182 | 1.82314 | 2.22 |
| 15 | 0.00286 | 2.85859 | 1.33 |
| 20 | 0.00441 | 4.41406 | 0.82 |
| 25 | 0.00674 | 6.73514 | N/A |

### 2.3 Curve fitting and shape analysis

The shape of the lipid-detergent complexes was deduced following the detailed method of Mazer and co-workers to characterize mixed micelle formation in bile salt-lecithin solutions. This method was used by Glover and co-workers to deduce the discoidal shape of DMPC-DHPC bicelles [4, 19]. The lipid-detergent solutions were treated as monodisperse and non-interacting where the mean scattering intensity, *I*, is given by the following equation:

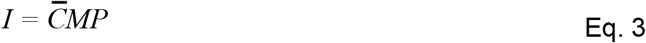

where *C* is the concentration of the lipid-detergent *(w/w), M* is the molecular weight of the lipid-detergent complex, and *P* describes the scattering form factor. The quantity *I/C* is proportional to *MP*. Using this approach, the product *MP* will have different values depending on the shape of the lipid detergent complex. It follows that the quantity *I/C*, which is equivalent to *MP*, can be measured experimentally for DMPC-C_8_E_5_ complexes and the shape deduced from a semi-log plot of the normalized *I/C* versus *Rh*. The dependence of *I/C* on *Rh* was fitted to one of three curves that were generated for complexes with a spherical shape (Figure 4 - solid line), disk shape (Figure 4 - dashed line), and a rod shape (Figure 4 - dotted line).

**Figure 4.**
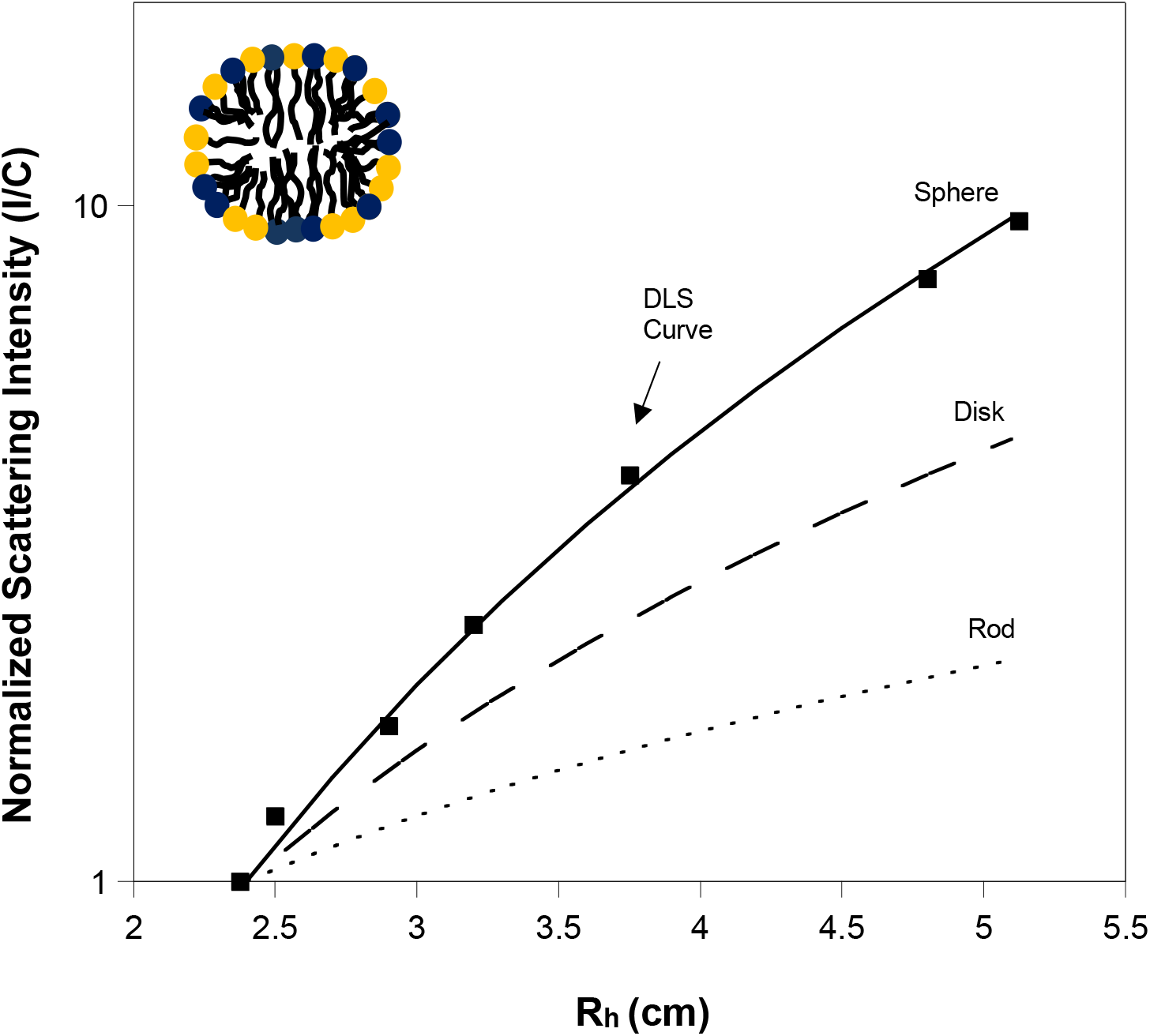
Semi-log plot of normalized scattering intensity versus radius of hydration, R_h_. The solid line represents the model of a spherical particle, the dashed line represents the model of a disk, the dotted line represents the model of a rod. The dotted-square line represents the DLS data. The DLS data fit the curve for the sphere indicating the shape of the DMPC-C_8_E_5_ complexes favor spheres over discoidal-shaped complexes at ***q*** ≤ 0.5.

## 2. Results and Discussion

### 3.1 Sedimentation equilibrium

#### Density matching of DMPC-C_8_E_5_ complexes

All sedimentation equilibrium experiments were performed at 25 °C using a Beckman XL-A analytical ultracentrifuge. To follow the sedimentation of the lipid complexes using absorbance optics, an NBD-labeled phospholipid probe was incorporated in a trace amount, 1:10,000 mole ratio of NBD-PE to DMPC.

Sedimentation equilibrium data was collected for DMPC-C_8_E_5_ lipid-detergent complexes at a ***q***-value of 0.25 and a total lipid concentration of 5% (w/w). A total of nine samples were prepared at D_2_O concentrations of 0.00, 3.25, 6.50, 9.75, 13.00, 16.25, 19.50, 22.75, and 26.00% D_2_O (v/v). Sedimentation profiles were generated for all nine samples at rotor speeds of 15,000 rpm (18,112 x g), 20,000 rpm (32,198 x g), 25,000 rpm (50,310 x g) and 30,000 rpm (72,446 x g). The sedimentation profiles were mathematically treated using the Lamm equation (Eq. 1) and the effective molecular weight, M_eff_, was calculated for each sample at each D_2_O concentration. M_eff_ was plotted against the percentage D_2_O concentration to determine how much was required to density match the DMPC-C_8_E_5_ or reach M_eff_ = 0. Figure 5 shows that approximately 18.4% D_2_O (v/v) is required to be matched, much lower than the 71.7% required to match DMPC-DHPC bicelles [2]. High quantities of density modifiers introduce the risk of generating a concentration gradient of the density modifier at high rotor speeds, which are commonly required when sedimenting small proteins. Thus, DMPC-C_8_E_5_ lipid-detergent complexes are advantageous over DMPC-DHPC bicelles for this reason.

**Figure 5.**
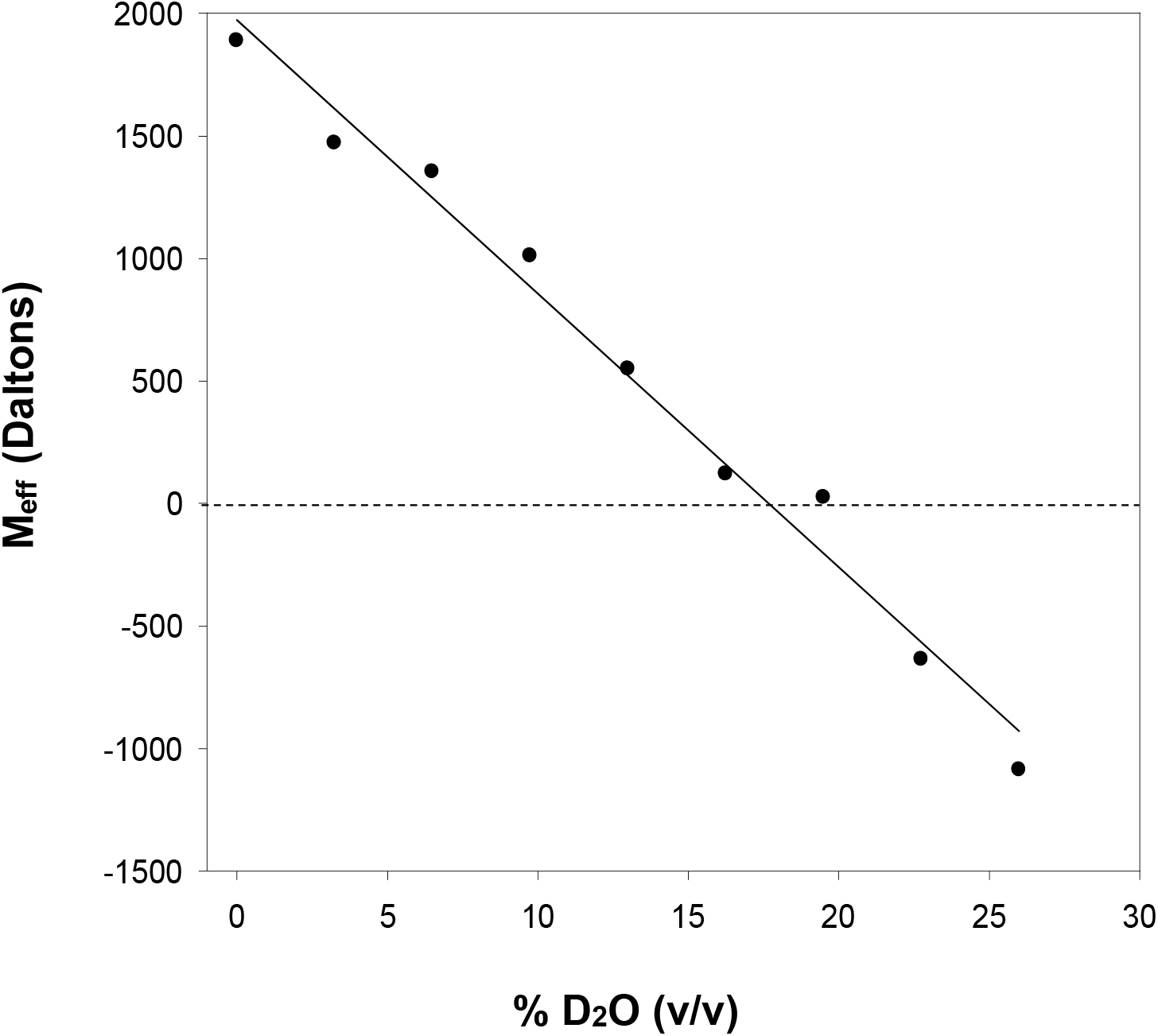
M_eff_ versus D_2_O concentration. Approximately 18.4% D_2_O is required to density match DMPC-C_8_E_5_ lipid-detergent complexes prepared at a ***q*** = 0.25.

### 3.2 Analysis of DMPC in bicelles and C_8_E_5_ complexes

To probe the lipid environment of DMPC in both DMPC-DHPC bicelles and C_8_E_5_ complexes, ^31^P-phosphorus NMR was used following the method of Glover and co-workers [4]. We reasoned that if DMPC assumed a similar arrangement in DMPC-C_8_E_5_ complexes then it follows that the chemical shifts would be the same or very similar to that of DMPC-DHPC bicelles. Figure 6 shows an overlay of both spectra at 25 °C and 37 °C (Figures 6A and 6B, respectively). We can see from these spectra that the chemical shift of DMPC is very similar to its chemical shift in DMPC-DHPC bicelles with only a slight shift. These experiments were run at both 25 °C and 37 °C to determine if there were any temperature-related effects, and although at higher temperatures this shift in the signal becomes more pronounced, there is little indication that temperature largely influences the lipid environment of DMPC in either case. We would predict DMPC to have a largely different chemical shift compared to DMPC-DHPC bicelles were it not associated with C_8_E_5_ in a bicelle-like environment, though it is unclear what the magnitude of that shift would be. The close similarity of the chemical shifts suggests that DMPC occupies a bilayer region in both bicelles and C_8_E_5_ complexes at ***q*** = 0.5.

**Figure 6.**
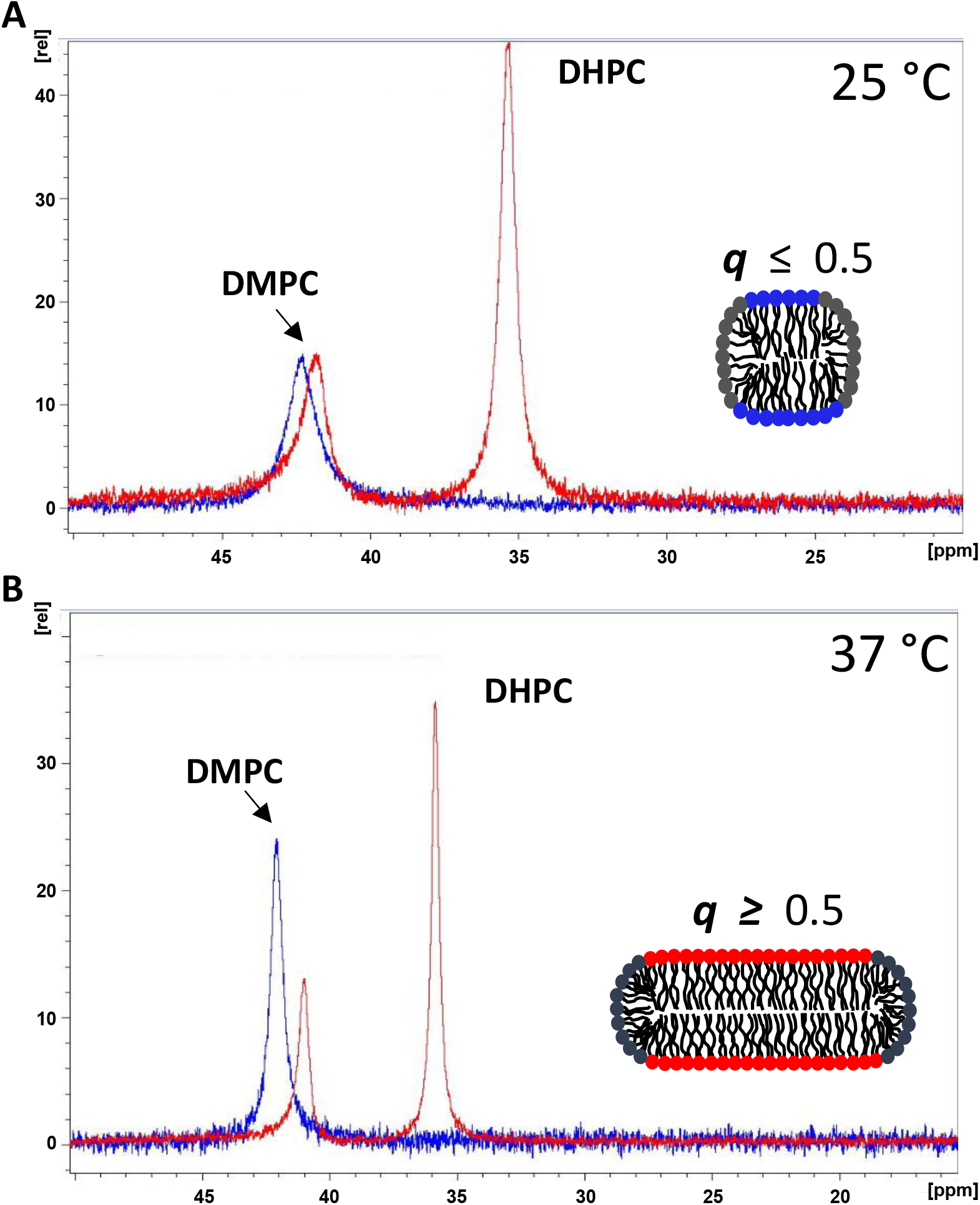
^31^P-phosphorus NMR spectra of DMPC in C_8_E_5_ lipid complexes (blue) and bicelles (red) at 25 ºC (A) and 37 ºC (B). DMPC-DHPC bicelles and DMPC-C_8_E_5_ lipid complexes were prepared at a ***q*** of 0.5. The chemical shift of DHPC is approximately 35.1 ppm in bicelles at both 25 and 37 ºC. The chemical shift of DMPC appears at 41.0 ppm at 37 ºC and 42.0 ppm at 25 ºC in bicelles. In C_8_E_5_ complexes DMPC appears at 42.5 ppm at both 37 and 25 ºC. The inset in each panels shows a schematic visualization of how small bicelles, ***q*** ≤ 0.5 (top), might look with a spherical morphology and a narrow bilayered region compared with larger bicelles, ***q*** ≥ 0.5 (bottom), with a broader planar region.

### 3.3 Dynamic light scattering to evaluate complex size and shape

To investigate the shape morphology of DMPC-C_8_E_5_ complexes, light scattering studies were undertaken using the method of Mazer and Glover [4, 19]. Samples were prepared at a ***q***-value of 0.25 at various % total lipid-detergent compositions ranging from 1% up to 25% (w/w). At a total lipid-detergent concentration of 25% (w/w), dynamic light scattering data could not be acquired because the lower limit of size detection had been reached. This trend in decreasing complex size could be seen across the different samples (Figure 3). One plausible explanation for this is that the concentration of freely available C_8_E_5_ in solution increases as the total lipid-detergent concentration goes up. When this happens, the resulting size of the complexes decreases because there is more freely available C_8_E_5_ in solution that can associate with DMPC to form smaller complexes.

From the Stoke’s-Einstein equation and the decay of the autocorrelation function, the hydrodynamic radius, R_h_, of the DMPC-C_8_E_5_ complexes was determined for each lipid-detergent sample and plotted versus ***q***-value. Figure 2 shows the size of the complexes, R_h_, increases with respect to ***q***. A similar increase in the hydrodynamic radius was reported for DMPC-DHPC bicelles, which initially supported our hypothesis that DMPC-C_8_E_5_ lipid-detergent structures assume a bicellar-like arrangement. To further investigate the shape of these structures we followed the method of Mazer and co-workers which was later applied by Glover et al. in detail [4, 19]. Using this approach, three curves were generated. Each describes the behavior of model spherical complexes (solid line), discoidal complexes (dashed line), and rod-like complexes (dotted line) (Figure 4). The molecular weight, *M*, was calculated for each type of complex along with a unique form factor, *P*. The product of the molecular weight and the form factor, *MP*, is equivalent to the experimentally determined scattering intensity, *I*, divided by the lipid-detergent concentration, *C*, for the DMPC-C_8_E_5_ structures:

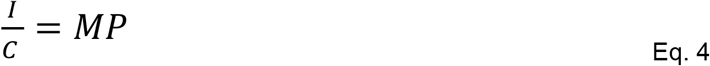

For the curves, the product of the molecular weight and form factor, *MP*, were calculated and plotted on a semi-log plot versus the hydrodynamic radius for spheres, disks, and rods. The range for the hydrodynamic radius was determined based on the measured hydrodynamic radius, R_h_, for the DMPC-C_8_E_5_ samples. These R_h_ values were substituted into the values of R_h_ for each of the models and the calculated scattering intensity was plotted in Figure 4. We chose to use dimensions of DMPC that corresponded to the previous findings by Glover and co-workers, which agreed with the calculations introduced by Vold and Prosser [4, 5, 20]. For the disk and rod-shaped models, the thickness, *t*, and the diameter, *d*, were kept constant at 5 nm, governed by the length of DMPC [4]. In these two models the radius of the disk, *r*, and the length, *L*, of the rod-shaped complexes were adjusted to the measured values of the radius for the DMPC-C_8_E_5_ complexes. For the spherical model, the radius, *a*, was left to float between values of 1 and 10. The data were normalized to the smallest value and ten points were obtained for each curve.

For DMPC-C_8_E_5_ the scattering intensity, *I*, was divided by the total lipid-detergent concentration, *C*, for each ***q***-value. The data were normalized to the lowest scattering intensity and plotted against the measured hydrodynamic radius, *Rh*. Our analysis revealed that when the normalized scattered light intensity was plotted on a semi-log plot versus the hydrodynamic radius for DMPC-C_8_E_5_, the resulting curve overlays remarkably well with the curve for a spherical micellar complex. A result that is more consistent with a micellar-like complex and not a disk-shaped complex. All C_8_E_5_ lipid samples were kept at a ***q*** ≤ 0.5 due to observed phase changes and reported changes in solution viscosity at higher ***q*** (Table 1). Viscosity also increased substantially when ***q*** was held constant at 0.25 and the total lipid concentration, [DMPC + C_8_E_5_], was varied from 1 to 25% (w/w) (Table 2).

DHPC and C_8_E_5_ have very different reported CMC values at approximately 16 mM and 8 mM, respectively [14, 21, 22]. The differences in CMC may account for the viscosity differences observed and measured for DMPC-C_8_E_5_ at higher ***q***. The lower CMC of C_8_E_5_ means it more readily forms micelles in solution. This means less free C_8_E_5_ is available to associate with DMPC potentially leading to a much higher ***q***_eff_ for the DMPC-C_8_E_5_ lipid complexes. For bicelles at high ***q***, ≥ 3.0, they assume more gel-like physical characteristics, which can be directly attributed to a lower availability of free DHPC in solution (see Eq. 2) [23]. All samples prepared with C_8_E_5_ in this study were carried out at concentrations above the CMC.

## 3. Conclusions

In this study a new lipid bicelle system was prepared using the phospholipid, DMPC and detergent, C_8_E_5_. Two important parameters that are used to describe bicelles were varied and the biophysical properties of the resulting complexes were evaluated; the ***q***-value and the total lipid concentration (% w/w) defined as the sum of the concentrations of DMPC and C_8_E_5_, [DMPC + C_8_E_5_]. The properties of the lipid-detergent complexes were characterized and found to assume a more sphere-like structure based on modeling studies using light scattering rather than disc-shaped complexes adopted by DMPC-DHPC bicelles. Further, the hydrodynamic radius, R_h_, of the C_8_E_5_ complexes at ***q*** = 0.5 was reported to be approximately 5.25 nm, which is more consistent with what we expect for detergent micelle complexes rather than bicelles, which tend to be larger at a similar ***q***. DMPC-C_8_E_5_ complexes were density matched in the analytical ultracentrifuge suggesting that they can be used for sedimentation equilibrium analysis aimed at studying membrane protein oligomerization. The significant increase in solution viscosity at ***q***-values greater than 0.5 suggests a “solution dynamic range” that is below that for DMPC-DHPC bicelles. Though we have used parameters typically reserved to describe the properties of DMPC-DHPC bicelles, it is important to note that this new system may not behave in the same way and necessitates further characterization of its physical properties expanding into a broader range of ***q***-values. Additionally, an investigation into their adaptability to NMR analysis of membrane proteins will be beneficial for structural studies.

## Abbreviations

DMPC: 1,2-dimyristoyl-*sn*-glycero-3-phosphocholine
DHPC: 1,2-dihexanoyl-*sn*-glycero-3-phosphocholine
C_8_E_5_: n-octylpentaoxyethylene
NBD-DMPE: (1,2-dimyristoyl-*sn*-glycero-3-phosphoethanolamine-N-(7-nitro-2-1,3-benzoxadiazol-4-yl) (ammonium salt))
^31^P-NMR: phosphorus nuclear magnetic resonance spectroscopy
D_2_O: deuterium oxide
CHAPS: 3-[(3-cholamidopropyl)dimethylammonio]-1-propanesulfonate
CHAPSO: 3-([3-Cholamidopropyl]dimethylammonio)-2-hydroxy-1-propanesulfonate
HEPES: 2-[4-(2-hydroxyethyl)piperazin-1-yl]ethanesulfonic acid
DLS: dynamic light scattering

## 4. Acknowledgements

I would like to thank those who supported this work by providing access to instrumentation and data collection. I would also like to thank the reviewers, past and present, who took the time to critically read versions of the manuscript and offer valuable feedback. Lastly, thank you to the Chemistry Department at Southern Illinois University, Edwardsville.

